# Dual-Approach Co-expression Analysis Framework (D-CAF) Enables Identification of Novel Circadian Regulation From Multi-Omic Timeseries Data

**DOI:** 10.1101/2024.10.10.617622

**Authors:** Joshua Chuah, Carmalena Cordi, Juergen Hahn, Jennifer Hurley

**Author notes:** Contributing authors (Carmalena Cordi, Juergen Hahn).

## Abstract

The circadian clock is a central driver of many biological and behavioral processes, regulating the levels of many genes and proteins, termed clock controlled genes and proteins (CCGs/CCPs), to impart biological timing at the molecular level. While transcriptomic and proteomic data has been analyzed to find potential CCGs and CCPs, multi-omic modeling of circadian data, which has the potential to enhance the understanding of circadian control of biological timing, remains relatively rare due to several methodological hurdles. To address this gap, a Dual-approach Co-expression Analysis Framework (D-CAF) was created to perform perturbation-robust co-expression analysis on time-series measurements of both transcripts and proteins. Applying this D-CAF framework to previously gathered transcriptomic and proteomic data from mouse macrophages gathered over circadian time, we identified small, highly significant clusters of oscillating transcripts and proteins in the unweighted similarity matrices and larger, less significant clusters of of oscillating transcripts and proteins using the weighted similarity network. Functional enrichment analysis of these clusters identified novel immunological response pathways that appear to be under circadian control. Overall, our findings suggest that D-CAF is a tool that can be used by the circadian community to integrate multi-omic circadian data to improve our understanding of the mechanisms of circadian regulation of molecular processes.

## 1 Introduction

Circadian rhythms are molecular rhythms that cycle once approximately every 24 hours in order to time organismal physiology to the day/night cycle of earth ([1]). These rhythms occur broadly in organisms that live in the Photic Zone ([2]) and play a crucial role in regulating physiological and behavioral processes, such as the sleep-wake cycle, autophagy, immune system function, hormone production, and metabolism ([3], [4], [5], [6], [7]). Given their wide-ranging regulation of human physiology, disruption of circadian rhythms has the ability to enhance the rates of human diseases, such as Alzheimer’s disease, diabetes, cancer, and cardiovascular disease ([8], [9], [10], [11]).

Because of the significant effect circadian rhythms have over physiology, much research has been devoted to understanding the drivers of these rhythms. The molecular mechanism that generates circadian rhythms is referred to as the molecular circadian clock and is comprised of a transcription-translation negative feedback loop that coordinates the rhythmic transcription/expression of a host of genes and proteins (called clock controlled genes/proteins, CCGs/CCPs) ([12], [13]). It is these CCGs/CCPs that are responsible for driving the 24-hour biological phenotypes that are collectively referred to as circadian physiology. With CCG/CCPs numbering in the hundreds to thousands depending on cell type, omics level data is often collected to characterize the effect the molecular circadian clock has on the transcriptome and proteome in an attempt to identify the mechanisms that link circadian disruption to disease ([14], [15], [16], [17]).

While many investigations into circadian rhythms use single omics approaches, like transcriptomics or proteomics, there is a growing recognition of the importance of modeling cellular functions using multiple omics, or multi-omics, datasets ([18], [19]). Multi-omic approaches are crucial in understanding the regulation of circadian physiology as oscillations in one data set (e.g. transcription factors in the proteome) can affect oscillations in another data set (e.g. rhythmic levels of expression in the transcriptome), which multi-omics can help to explore. One analysis technique that has been used independently in both multi-omics analysis and circadian biology is co-expression analysis, which finds clusters of analytes with similar expression patterns to define regulatory relationships within and between pathways ([20], [21]). Co-expression network analysis represents large scale data as a collection of nodes and interactions between nodes, which provides researchers with the ability to study the structure of complex systems using omics level data, enabling the clustering and exploration of modules of genes, transcripts, or proteins ([22])([23]).

While co-expression may be highly useful in defining the network of the circadian clock, to the best of our knowledge, co-expression analyses that integrate multiple omics datasets of circadian data simultaneously have only been used in a limited fashion ([22]). This lack of multi-omic co-expression analyses of circadian data is likely due to a plethora of hurdles to multi-omic analysis, many of which involve modeling of the relationships between multi-omics data from time series data. In fact, while there are several techniques that exist for the integration of multiple omics datasets, few have been proven to be useful for longitudinal data ([24]). Those that can be applied to longitudinal data either involve data transformation or necessitate modeling step pre-integration ([25], [26], [27]). These limit the ability to perform co-expression analysis, as transformation of the original expression patterns leads to a loss of direct interpretability, making it challenging to assess the similarities of the original data over time.

One way to mitigate these hurdles is to implement a strategy known as early integration. Early integration is the simplest multi-omic data integration method and involves concatenating multiple omics datasets together. However, this method is not frequently used as raw transcript and protein measurements can vary by orders of magnitude ([28]). Our suite of software packages, the Pipeline for Amplitude Integration of Circadian Exploration (PAICE) suite, including the Extended Circadian Harmonic Oscillator (ECHO) program, allows us to overcome this as it fits relative expression curves to longitudinal biological data that are comparable between data sets ([29]). Given that ECHO allows for early integration, we hypothesize that we are able to perform true multi-omics co-expression analysis on circadian multi-omics data.

Therefore, the aim of this study was to 1) develop a framework to perform co-expression analysis of circadian multi-omic data and 2) use this framework to identify circadian rhythms and rhythm-regulated functions in mouse macrophages. To do so, circadian transcriptomic and proteomic data previously collected from mouse macrophages were first preprocessed to enable the comparison of transcript and protein time-series using our ECHO program. Subsequently, multiple network analysis models were applied to 1) concurrently compute rhythms with the most similarities to known circadian rhythms and 2) create clusters that are then utilized to construct protein-protein interaction networks. The clusters we identified using this framework showed that circadian rhythms drive aspects of the immune system and ribosomal RNA processes in mouse macrophages. We found our co-expression framework to be useful in identifying densely connected, perturbation-robust communities, and larger, biologically relevant clusters. Moreover, we have designed a framework for others to apply this approach to their own longitudinal multi-omic datasets. In total, our Dual-approach Co-expression analysis Framework (D-CAF) builds a foundation that can be used broadly to analyze circadian multi-omic data.

## 2 Materials and Methods

A description of the application of the Dual-approach Co-expression analysis Framework (D-CAF) to a multi-omic (transcriptomic and proteomic) circadian time-series dataset is given below. A flowchart representing the overall steps in the framework can be found in Figure 1. In short, the framework can be broken down into a data pre-processing step, where early integration of the transcriptomic and proteomic datasets are performed, a modeling step, where models are generated to group rhythms with similar expression patterns over time, and a validation step, where the models are validated using various methods.

**Fig. 1.**
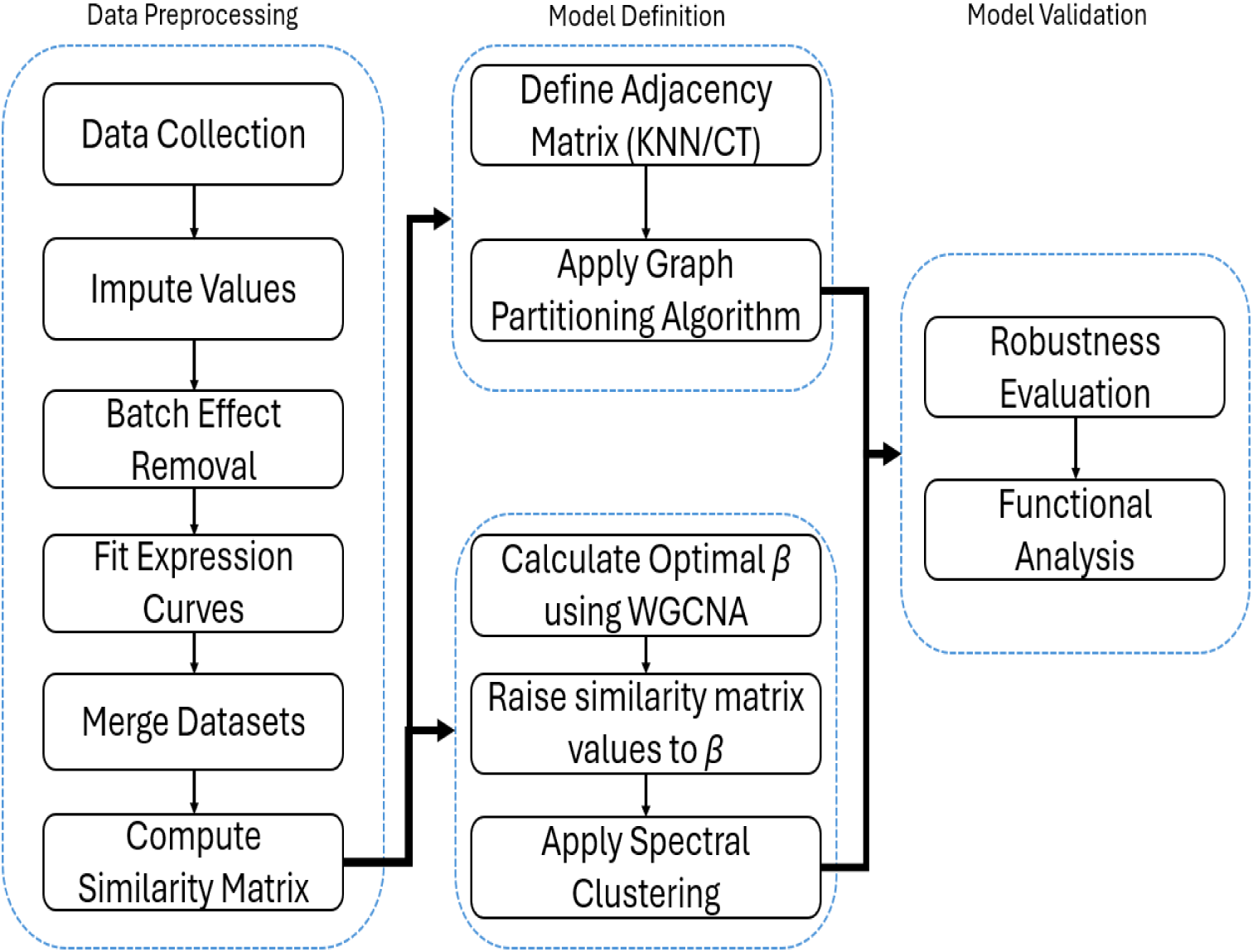
A flowchart for the general D-CAF workflow. Data was first preprocessed to integrate the individual omics datasets, after which both weighted and unweighted models were generated, and finally validation was run on the optimal models.

### 2.1 Data Collection

Data from [30] were used as source data as model data for our framework development. In this previously published data set, macrophages were collected from the bone marrow of Per2::Luc or Per1/Per2 knockout mice aged 3-6 months old. Transcriptomic measurements (RNA-sequencing data) were collected every two hours over a 48 hour window across three (independent) biological replicates. RNA-seq counts were computed with HTseq, and the raw counts data were normalized to transcripts per million. Proteomic measurements were also collected every two hours over a 48 hour window across three biological replicates. Protein concentrations were measured via multiplex tandem mass tag mass spectrometry. Further technical details on this data set and data collection can be found in the source paper for the data ([30]).

### 2.2 Data Preprocessing

Early integration of the proteomic and transcriptomic datasets were performed to merge both datasets into one multi-omic dataset. For early integration, both data sets had to be transformed and normalized such that there was minimal heterogeneity between data sets. Since raw expression values of transcripts and proteins can vary by factors of hundreds to thousands, our approach consisted of employing the LIMBR and ECHO software packages to create comparable expression curves. LIMBR first performed missing value imputation and batch effect removal of each individual replicate for both the proteomic and the transcriptomic datasets individually. After LIMBR, the data was processed through the ECHO software package in order to fit a smoothed relative expression curve for all three biological replicates ([29],[31]).

#### 2.2.1 Missing Value Imputation and Batch Normalization

To impute missing data and batch normalize the individual omics datasets, values in the transcriptomic (transcripts per million) and proteomic (relative peptide concentration) datasets were log2 normalized to reduce the skew and range of values at each time point while still remaining representative of the proportional differences at each time point. Next, unique peptides or reads from a given peptide or transcript respectively were summed into one measurement, so each row in the dataset was a unique peptide or transcript at a given time. Imputation was then performed using the K-nearest neighbors algorithm with K = 10, with only rows with less than 30% values missing imputed to limit over-imputation. Next, batch effect removal was performed to center the measurements around the same distribution to eliminate noise using surrogate variable analysis. Once the data was batch effect normalized, measurements from different peptides belonging to the same protein were averaged together. Further information on this process can be found in the source paper ([31]).

#### 2.2.2 Expression Curve Fitting

To create a smoothed relative expression curve for each protein or transcript, we employed our ECHO program ([29]). ECHO analysis normalizes biological replicates such that the transcript or protein mean is 0 and the row standard deviation is 1. This normalized data is then fitted to a regression model, resulting in a relative expression profile for each transcript and protein that is a weighted combination of the data from the three replicates. ECHO also computes several parameters of the curve, such as the initial amplitude, amplitude change coefficient, rhythm period, phase shift, and the goodness of fit p-value, which were all useful properties to verify the similarity of two rhythms. Further information on this process can be found in the source paper ([29]).

Using the combined LIMBR and ECHO approach, the rhythms of 36,000 transcripts and 6,000 proteins were fitted. To ensure that the vast majority of rhythms analyzed were representative of the original data, only transcripts or proteins with a B-H adjusted p-value less than 0.01 were considered for further analysis ([32]). After B-H score filtering, there were 10092 transcripts and 1986 proteins left for analysis. Since relative expression curves of both the transcriptomic and proteomic data were generated using the same LIMBR + ECHO workflow, this data was merged into one multi-omic dataset consisting of 12,078 rhythms. This resulted in a final dataset of 12,078 rhythms (rows) and 24 columns (time points).

### 2.3 Network Analysis

Once we created our merged multi-omic dataset, we next sought to characterize similarities between the expression profiles of transcripts and proteins in the multi-omic dataset using network analysis. To build a network, first an *N x N* similarity matrix, where *N* is the total number of analytes (12,078) in the dataset, was constructed ([33]). Each element of this matrix, s[i,j] corresponded to the similarity between analytes *i* and *j*, where *i* and *j* were between 1 and 12,078. The similarity metric used in this study was the Pearson correlation between each pair of rhythms. As such, the similarity between rhythm i and rhythm j, e.g., s[i,j], was always between 1 (perfectly in-phase, overlapping) and -1 (perfectly antiphase). Networks were then defined by converting the similarity matrices into adjacency matrices using a dual-approach network analysis consisting of an unweighted network and a weighted network. Unweighted networks were represented by discrete adjacency matrices where each value in the similarity matrix was either converted to a 1, indicating two rhythms are “connected” in the network, or a 0, indicating no connection ([34]). These networks are computationally efficient and effective in modeling specific outcomes or circumstances due to their binary nature. However, they may lack information about the resulting model. Weighted analysis, conversely, uses a continuous adjacency matrix that involves transforming the values in the similarity matrix such that larger correlation values have a larger separation from smaller correlation values ([33]). In this way, weighted networks are more accurately able to capture groups of highly correlated rhythms, but they are computationally more expensive. Given their strengths and weaknesses, we used both approaches to capture local similarities between rhythms through unweighted network analysis and larger functional groups through weighted network analysis.

### 2.4 Unweighted Network Analysis

In our framework, unweighted network models were developed to find small communities of similar rhythms that show related patterns over time as compared to known core clock genes and proteins. To create these unweighted networks, we computed adjacency matrices using two methods: K-nearest neighbors (KNN) and correlation thresholding (CT). These methods are defined further below. Groups of similar expression patterns were then computed via several graph partitioning algorithms. These models were then validated with a robustness evaluation technique to determine if the computed communities are resilient to noise, e.g., if the similarity between two rhythms is stable in the presence of noise. The optimal unweighted network model was then selected and analyzed based on an assessment of all computed models’ robustness.

#### 2.4.1 Correlation Thresholding

Correlation thresholding (CT) is a method to generate adjacency matrices that is often used in biological co-expression analyses as the results are relatively easy to interpret due to the use of a simple threshold of the similarity matrix (correlation matrix) ([35]). Correlations above a threshold are marked as adjacent (1), while the rest are marked as non-adjacent (0). In circadian studies, the absolute value of the correlation in CT is typically used to group together in-phase and anti-phase rhythms. However, as the goal of this study is to identify similar expression patterns among circadian rhythms, adjacency matrices were built off a signed correlation matrix instead. In addition, as there are relatively few time points, we necessitated a high correlation between rhythms to consider two rhythms sufficiently similar to be considered related.

To determine a suitable correlation threshold, the percent of shared nearest neighbors was plotted against the correlation between circadian rhythms. If there were very few shared nearest neighbors at a correlation threshold, then this threshold would likely be too low to consider two rhythms co-expressed. To set the parameters for this procedure, we found the average number of overlapping nearest neighbors at several correlation levels among transcripts and proteins with a circadian period (Figure 3). In general, there were very few, if any pairs of analytes with overlapping neighbors when the correlation between those analytes was less than 0.9. This demonstrated that setting a CT above 0.9 was an appropriate threshold for network analysis and the minimum CT used to create adjacency matrices was therefore set to 0.9. To garner a better understanding of the correlation, three adjacency matrices were computed in this work, using a CT of 0.9, 0.95, and 0.99.

#### 2.4.2 K-Nearest Neighbors

K-Nearest Neighbors (KNN) is a widely used approach to identify similarities in biological data that, as opposed to CT, ranks each correlation to a given rhythm, sets the highest K to 1, and sets all other relationships to 0 ([36], [37]). While this can mean that some barely correlated rhythms are marked as related, KNN benefits from being able to use directional relationships (e.g., *i* can be the nearest neighbor of *j*, but *j* does not have to be the nearest neighbor of *i*).

To determine the parameters of the adjacency matrix using KNN, we found the apt values of K by calculating the standard deviation of the period, phase shift, and amplitude change coefficient among the K nearest neighbors, ranging from 10 to 150 nearest neighbors, in intervals of 10. Based on the low standard deviation of parameters, K values of 20, 40, 80 and 100 were found to be appropriate and non-redundant and were therefore used to construct the adjacency matrices (Figure 3B, C, and D).

#### 2.4.3 Model Generation

Using both a CT and KNN approach, we build our D-CAF framework to generate models that identify groups of similarly expressed rhythms over time by partitioning a co-expression network into communities. These models then needed to be validated by evaluating their robustness to noise. To complete the partitioning step, graph partitioning (GP) algorithms, also known as community detection (CD) algorithms, were applied to partition the network into densely connected communities (clusters) of rhythms ([38]). In this framework, four CD algorithms were used: Leiden, RB Potts (RBP), Asymptotic Surprise (AS), and Significance ([39], [40], [41], [42], [43]). Each algorithm was applied to all 7 (3 CT, 4 KNN) computed unweighted networks (defined by adjacency matrices). In total, 28 models were generated in this study by applying each of the 4 community detection algorithms to each of the 7 adjacency matrices. A graphical representation of this process is shown in Figure 2. These models partitioned sets of rhythms into communities, resulting in a set of cluster labels for each model that indicated, for each rhythm, which cluster that rhythm belongs to. From these 28 models, the optimal model was then selected via robustness evaluation, e.g. testing how a model’s cluster labels change when noise is introduced to the data.

**Fig. 2.**
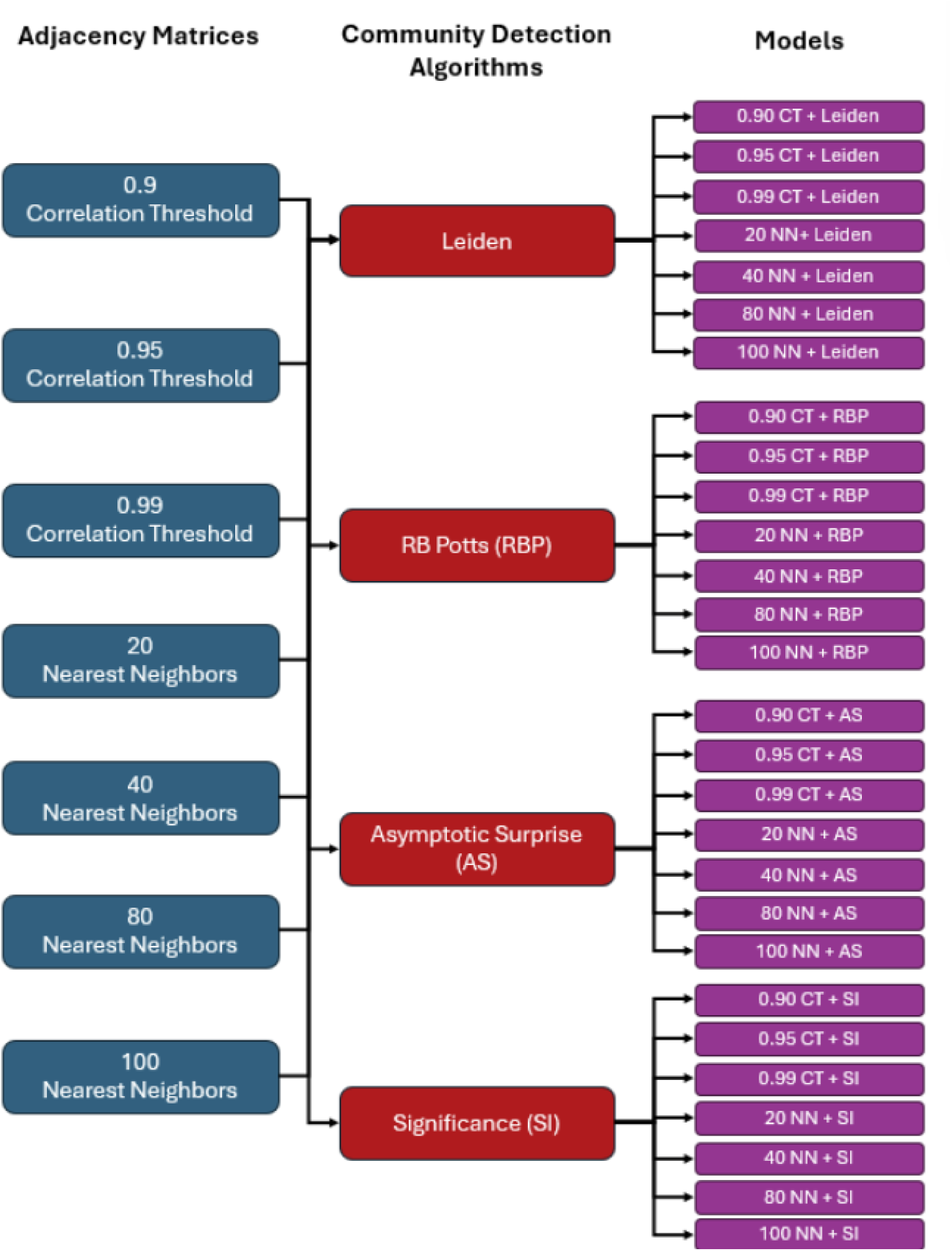
A graphical representation of the origin of the 28 unweighted models used in this study. 7 adjacency matrices were clustered using 4 community detection algorithms to create a total of 28 distinct models.

As the purpose of this study was to specifically find circadian clusters, the clusters produced by the models were further filtered by identifying clusters with an average circadian period (20-28 hours) and small standard deviation in period. The overall goal was to develop clusters with high within-cluster similarity and low between-cluster similarity. To this end, standard deviation in the ECHO-derived oscillation parameters (phase shift and period) were computed to determine the similarity of the rhythms contained within a cluster. If the standard deviation of these parameters was low (e.g. ¡ 2 hours), then the rhythms within a cluster were considered to be highly similar. Furthermore, the modularity of these clusters were computed to ensure the overall partitioned graph had many connections within each cluster and few connections between clusters.

### 2.5 Weighted Network Analysis

While unweighted community detection finds small communities of rhythms, weighted network analysis can identify large functional groups of co-expressed rhythms. To do so, weighted networks were generated by weighting factor *β* of the similarity matrices computed by the Weighted Gene Co-Expression Analysis package (WGCNA) based on the truncated *R*^2^ value ([33]). The weighted similarity matrix was then generated by raising each element of the original similarity matrix to a power *β*. By doing this, high correlations remained high and low correlations converged towards zero. Spectral clustering was applied to create clusters from this weighted similarity matrix, primarily because it 1) was faster for continuous similarity matrices and 2) allowed for the specification of the number of clusters to compute to avoid clusters that are too small. 30 clusters were chosen from the WGCNA analysis as the standard deviation of the within-cluster period plateaued for greater amounts of clusters, indicating larger numbers of clusters did not create significantly more tight-knit clusters (Figure S1.1). The robustness of the resulting model was then computed using the same procedure as the unweighted models to determine the reliability of the model.

### 2.6 Functional Enrichment Analysis

Circadian clusters were identified by selecting only clusters with average periods that were approximately 24 hours (between 20-28 hours). Biological functions in clusters were analyzed via protein-protein interaction (PPI) networks modeled by StringDB ([44]). Since the number of transcripts in this data outnumbered the amount of proteins, there are many instances where each transcript does not have its corresponding protein in the dataset. As such, analysis via methods that would analyze transcript-protein pairs such as Reactome or PathviewDB analysis prove difficult with this dataset. Therefore, PPI networks were chosen for biological analysis of clusters, as it enables comparison of these transcripts with the proteins that were detected. By inputting the members of each cluster, StringDB outputs a protein-protein interaction network where nodes are analytes and edges indicated interactions. To analyze this network, the PPI enrichment p-value was measured to determine if the number of connections within the cluster was significantly greater than the number of connections that would be expected in a random clustering of analytes. If there were significantly more (p *<* 0.05) edges than expected in the PPI network, then the clusters identified had some functional significance. Finally, gene ontology and reaction pathway analysis was performed to determine potential pathways and functions that the molecules in each cluster were related to.

### 2.7 Robustness Evaluation

After defining both the weighted and unweighted models, the models then had to be validated to ensure quality. To do so, model robustness was evaluated by determining how stable a model’s clustering output, referred to as *C*, was when noise was introduced to the data. This was implemented as biological data can be inherently noisy, and slightly different measurements in replicates could result in completely different networks and clustering outputs. If a community detection model is robust to noise, it means the relationship between the rhythms that were clustered are sufficiently strong to ignore small amounts of noise. To do so, noise was simulated by adding or subtracting a number randomly generated by a Gaussian distribution with mean = 0 and standard deviation = *σ* to each element in the original data matrix. The values of *σ* used in this study were 0.05 to 0.5 in increments of 0.05, with greater values of *σ* indicating greater severity of noise. This range was chosen to test how reliable a community detection model was with only a small amount of noise (0.05) and severe noise that would be expected to cause a decrease in robustness (0.5).

After applying Gaussian noise, the adjacency matrix was then re-created for each network and community detection was reapplied to generate *C\**, the recomputed cluster labels. The adjusted mutual information (AMI) score between *C* and *C\** was used to measure the difference between these original cluster labels and the recomputed cluster labels. When comparing two sets of cluster labels for a series of data points, we used an AMI score of 1 to represent total overlap, and an AMI score of 0 to represent total randomness ([45]). This was repeated 50 times for each model per *σ*. The average AMI score among these 50 repetitions was recorded as the model’s robustness to the severity of noise (*σ*). Models with greater AMI were deemed more robust to noise, and therefore were a higher quality model.

**Fig. 3.**
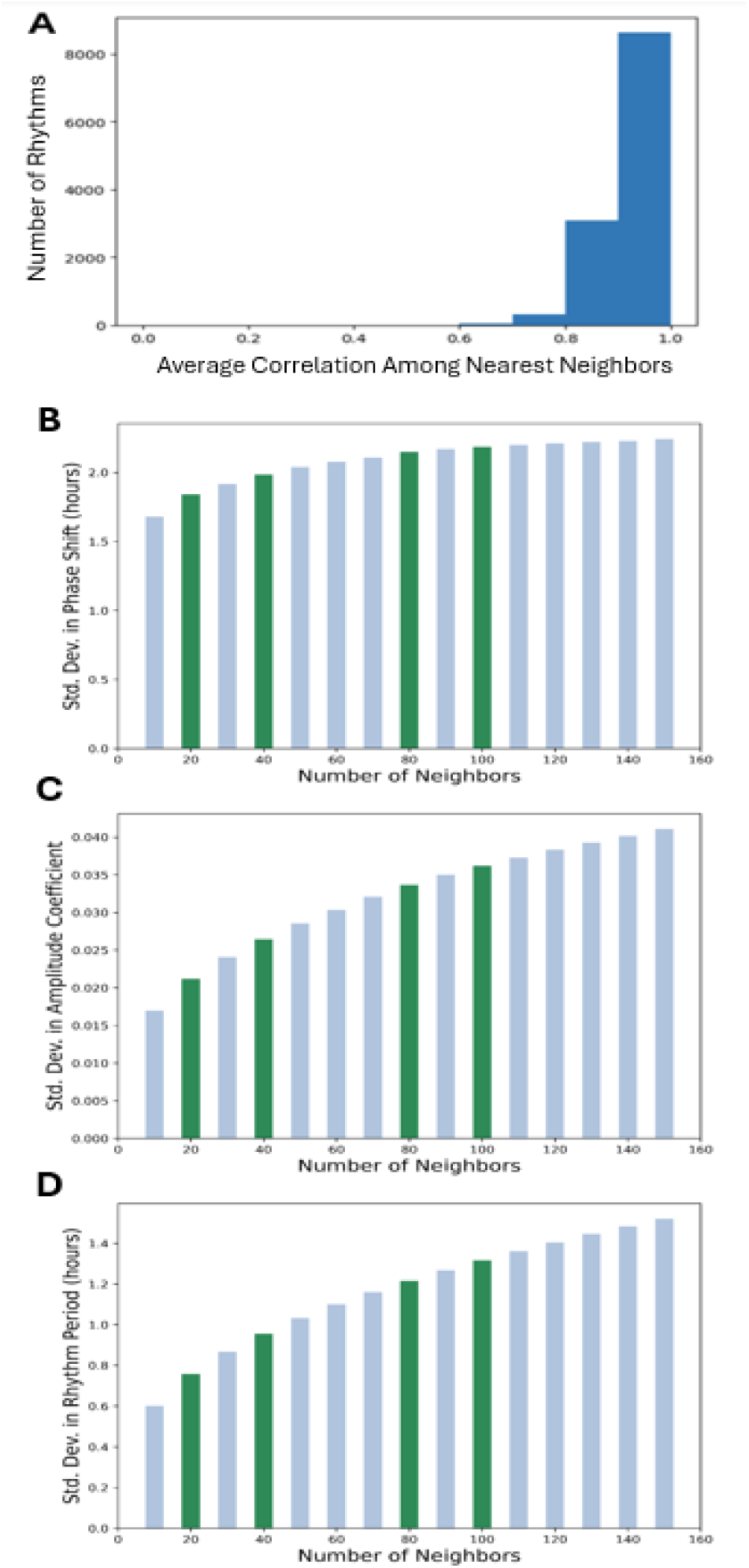
Parameter selection for adjacency matrices used in unweighted model construction. (A) shows the average correlation among each rhythm’s 100 nearest neighbors. (B), (C), and (D) show the average standard deviation in phase shift, period, and amplitude coefficient respectively among a model’s K-nearest neighbors. Bars in green indicate values of K selected to create adjacency matrices for unweighted model construction.

## 3 Results

### 3.1 Unweighted Network Analysis of Macrophage Transcripts and Proteins

Once we had a sound pipeline for D-CAF, we subjected our previously analyzed data from Collins et al. to D-CAF unweighted analysis, with the goal of identifying densely connected clusters of rhythmic elements that were also robust to perturbations ([30]). We measured the robustness of the original 28 candidate models (Figure 2) and computed the adjusted mutual information (AMI) score during repeated perturbation analysis. Heatmaps containing the average AMI score of each model when perturbed by small (*σ*=0.05) and severe (*σ* =0.5) noise are presented in Figure 4 (A and B). A higher AMI (close to 1) indicates a more stable model, whereas a lower AMI indicates that the model was not robust to noise, and is therefore not reliable ([46]).

**Fig. 4.**
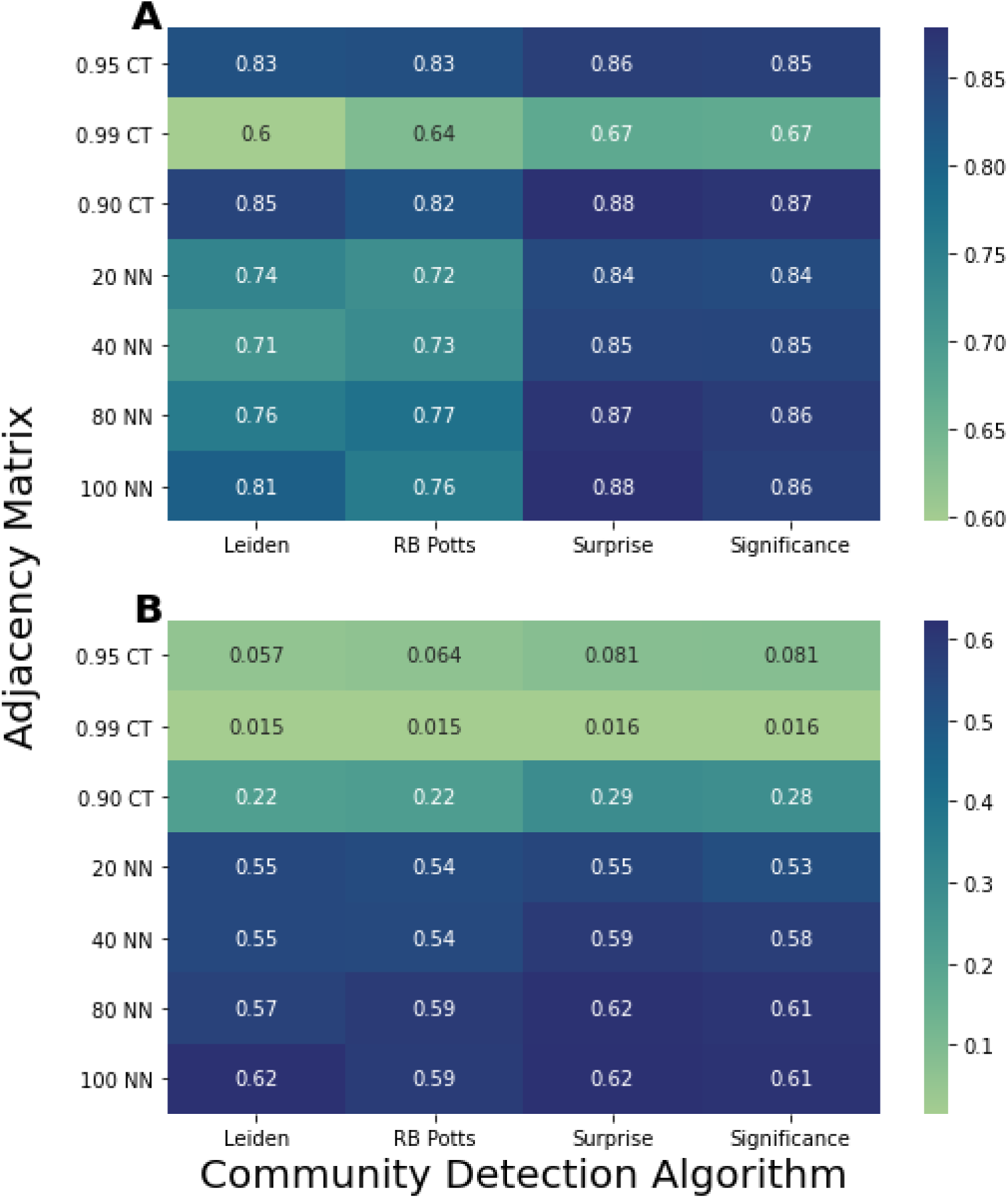
AMI scores identify optimal models to be used for Unweighted Network Analysis. Heatmaps displaying the adjusted mutual information (AMI) of clusters predicted by unweighted analysis when data was perturbed by Gaussian distributed noise with a standard deviation of 0.05 (A) and 0.5 (B). Values closer to 1 indicate higher similarity between the perturbed and original clusters and implies greater robustness.

At low noise severity (sigma = 0.05), we found the most robust models were those that were generated using the Asymptotic Surprise (AS) and Significance community detection algorithms. In general, the Leiden and RB Potts community detection algorithms were less robust than the AS and Significance community detection algorithms, likely because the AS and Significance algorithms are known to be able to be better at forming small communities, which is the goal of unweighted network analysis. In investigating the matrices, we found only the 0.99 CT Adjacency matrix was not as robust as all the other matrices. This is likely due to the conservative correlation threshold, where even slight perturbations would cause a rhythm to fall below the correlation threshold of 0.99. At severe noise (sigma = 0.5), we found the maximum AMI was achieved when the 80 and 100-NN matrices were paired with the AS community detection algorithm (Figure 4B). Interestingly, all CT-based clustering models generated low AMI scores when challenged with severe noise, indicating that these cluster models were not very reliable. This data reinforces the idea that candidate models should be evaluated at multiple severities of noise to ensure cluster model robustness when performing co-expression analysis.

Given all of the above, the 80-NN + AS model was selected and used for the unweighted co-expression analysis of the data from Collins et al. ([30]). The 80-NN + AS model partitioned the data into 84 clusters of rhythms. Of these, 18 were found to have a circadian period (i.e. the average period +/- standard deviation for these clusters was between 20-28 hours). Within each circadian cluster, the average standard deviation in period was 1.49 hours, indicating that the rhythms in each cluster oscillated at approximately the same frequency. Furthermore, the average standard deviation in phase shift within each cluster was 1.86 hours, indicating that rhythms in the each cluster shared the same circadian phase. A cluster-by-cluster breakdown of the cluster periods and phase shifts is presented in Table S1.1. The low standard deviation of rhythm parameters within each cluster demonstrated that the 80-NN + AS model successfully grouped similar rhythms without requiring explicit information about the rhythm parameters (e.g., phase shift and period).

We next computed the modularity score of the clusters generated by the 80-NN + AS model. The modularity of a cluster model score can fall between -1 and 1, and a score closer to 1 means there are many connections within a cluster and few connections between clusters (i.e., clusters are dense and distinct), which is optimal for a clustering model ([47]). Previous research has indicated that a modularity above 0.3 indicates that the structure of the network is not random ([48]). The computed modularity of the 80-NN + AS model used in this study was 0.640, demonstrating that the clusters found were both densely connected within each cluster and sparsely connected between clusters, further supporting that the 80-NN + AS model successfully clustered our data.

When analyzing what transcripts and proteins were identified within the clusters, we noted that transcripts/proteins that are a part of the core circadian transcription-translation negative feedback loop were found within clusters identified by the 80-NN + AS model that oscillated with a circadian period (Figure 5). For example, Per1 and Cry2, two genes that encode for components that regulate the core clock mechanism, were found in the cluster represented in Figure 5A, while Cry1 and Npas2 were found individually in other clusters (Figure 5B and C) ([12]). Notably, the cluster with Per1 and Cry2 also contained the immune proteins Tm9sf1, Ptcra, and NTAL, all genes/proteins not previously shown to have a relationship with the circadian clock or circadian output ([49], [50], [51]). These genes/proteins have a correlation score of 0.884, 0.995, and 0.676 with respect to Per1, indicating that the unweighted analysis was able to identify and cluster novel genes/proteins with known circadian clock genes. As such, the clusters generated by our unweighted analysis present novel avenues of investigation for circadian regulation.

**Fig. 5.**
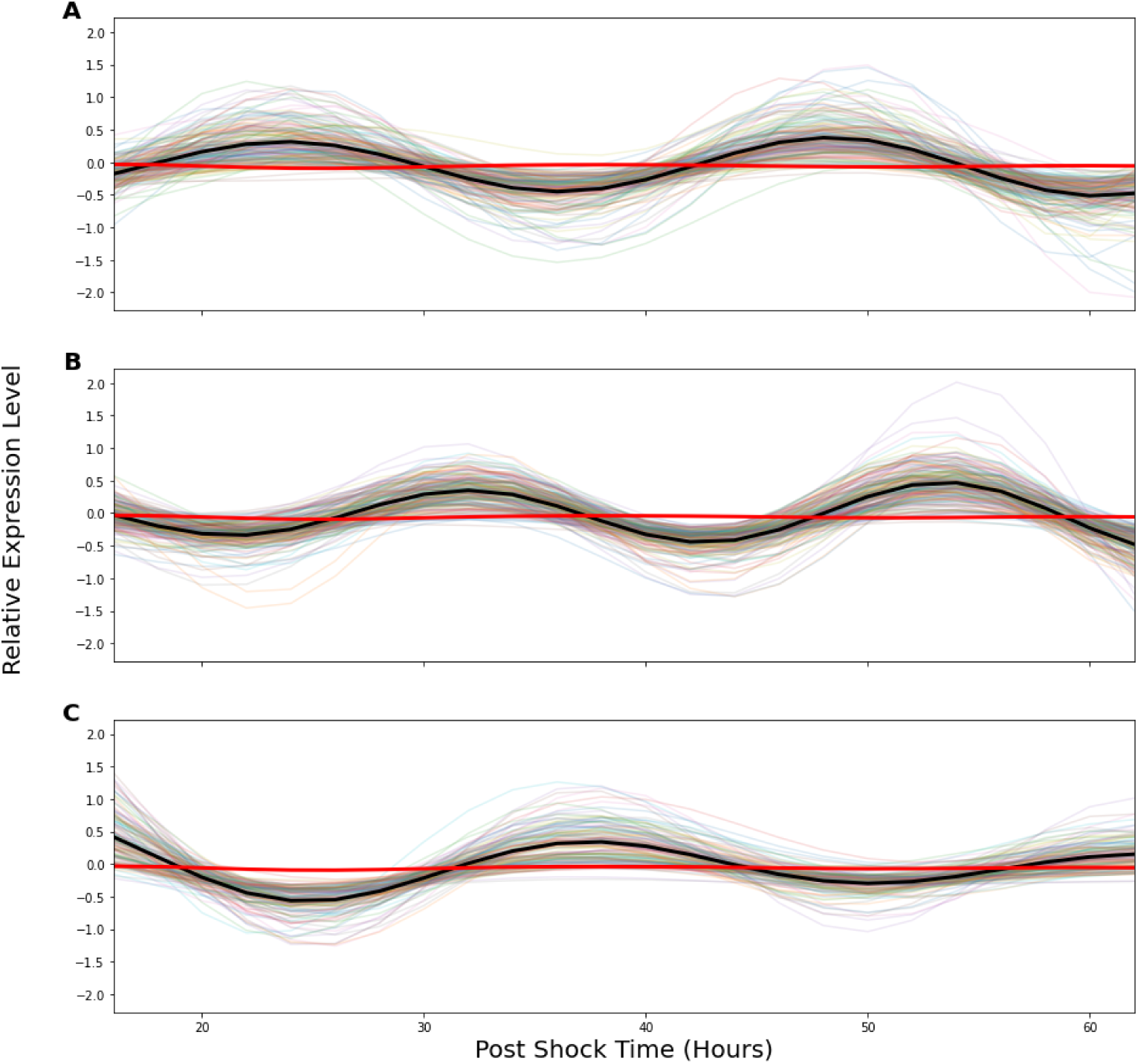
Expression profiles of circadian clusters identified by Unweighted Network Analysis demonstrate successful clustering. Each displayed cluster was found to have a circadian period, with the average expression profile for the cluster bolded in black. As a comparison, the average expression profile for the whole dataset is displayed bolded in red. The clusters contained 173 genes/proteins (A), 189 genes/proteins (B), and 137 genes/proteins (C).

### 3.2 Weighted Network Analysis of Macrophage Transcripts and Proteins

Although the small deviation in rhythm parameters from the unweighted network clusters indicated high similarity between individual rhythms within the same cluster, deriving global biological conclusions from these clusters was difficult as gene ontology or pathway analysis requires a larger sample size to find enriched functions. For this reason, we next employed the weighted network analysis function from D-CAF to analyze the data from Collins et al. ([30]) to develop clusters that could elucidate more broad, large-scale network information within the multi-omics dataset. To this end, fewer, larger clusters were generated by applying spectral clustering to the weighted network (SC + WN) model as described above. Of the 30 clusters identified, 5 were found to have a circadian period (i.e. the average period +/- standard deviation for these clusters was between 20-28 hours). The average standard deviation in period among these clusters was 2.28 hours, and the average standard deviation among the phase shifts was 1.95 hours. Although these standard deviations were significantly larger than the standard deviation of parameters among the circadian clusters generated by using an unweighted network approach, they were still close to the resolution of the time series used to generate the data (2 hours).

The SC + WN model was validated by repeatedly perturbing the model by adding Gaussian noise to the data, after which the AMI score was computed, with a larger *σ* indicating more perturbation to noise. As expected, the average AMI score of the SC + WN model decreased as noise decreased, but was similar to the AIM scores found in the unweighted network analysis (Figure 6). Overall, this indicated similar, albeit slightly less, robustness to noise in the weighted vs the unweighted analysis, and that the models generated by the SC + WN model were properly clustering patterns within the data.

**Fig. 6.**
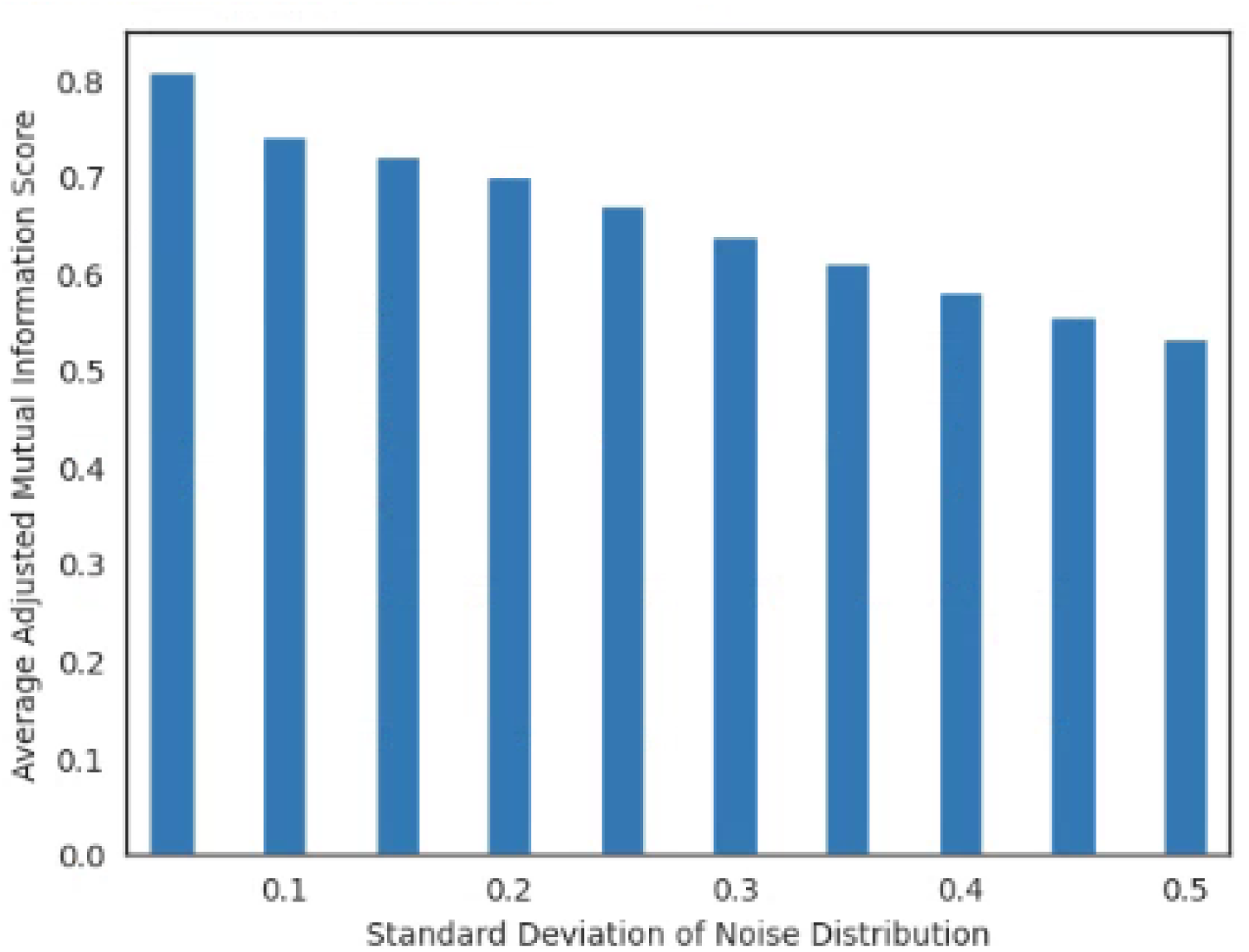
AMI scores for Weighted Network Analysis are similar to scores for the Unweighted Network Analysis. Higher AMI indicates higher overlap between the perturbed clustering and the original clustering. The severity of noise is represented by the standard deviation of the noise distribution. AMI for each severity of noise is averaged over 50 repetitions.

StringDB was used to find biological functions regulated by analytes with similar expression profiles to known clock transcripts and proteins. To this end, PPI enrichment networks were constructed from each of the SC + WN circadian clusters and we found that three of these networks had a PPI enrichment p-value less than 0.05, indicating that the cluster contained more biologically related analytes than would be expected in a random list of analytes. A list of p-values for each of the PPI networks computed from the SC + WN circadian clusters is available in Table S1.2. One such PPI network showed that rhythms from the IL-27 signaling pathway were contained in the same cluster, and therefore had highly similar expression patterns to known clock genes in mouse macrophages (Figure 7A). In addition to finding functionally related clusters, weighted clusters uncover more connections than the unweighted clusters (Figure 7B). When we created PPI enrichment networks from the unweighted and weighted modeled data, we found that, though the same module of IL-27 analytes were clustered with core clock genes, the weighted modeled data identified more auxiliary connections (Figure 7A and Figure 7B), demonstrating while both the unweighted and weighted models used in this study identify the same core information, the SC + WN model creates a more comprehensive functional network.

**Fig. 7.**
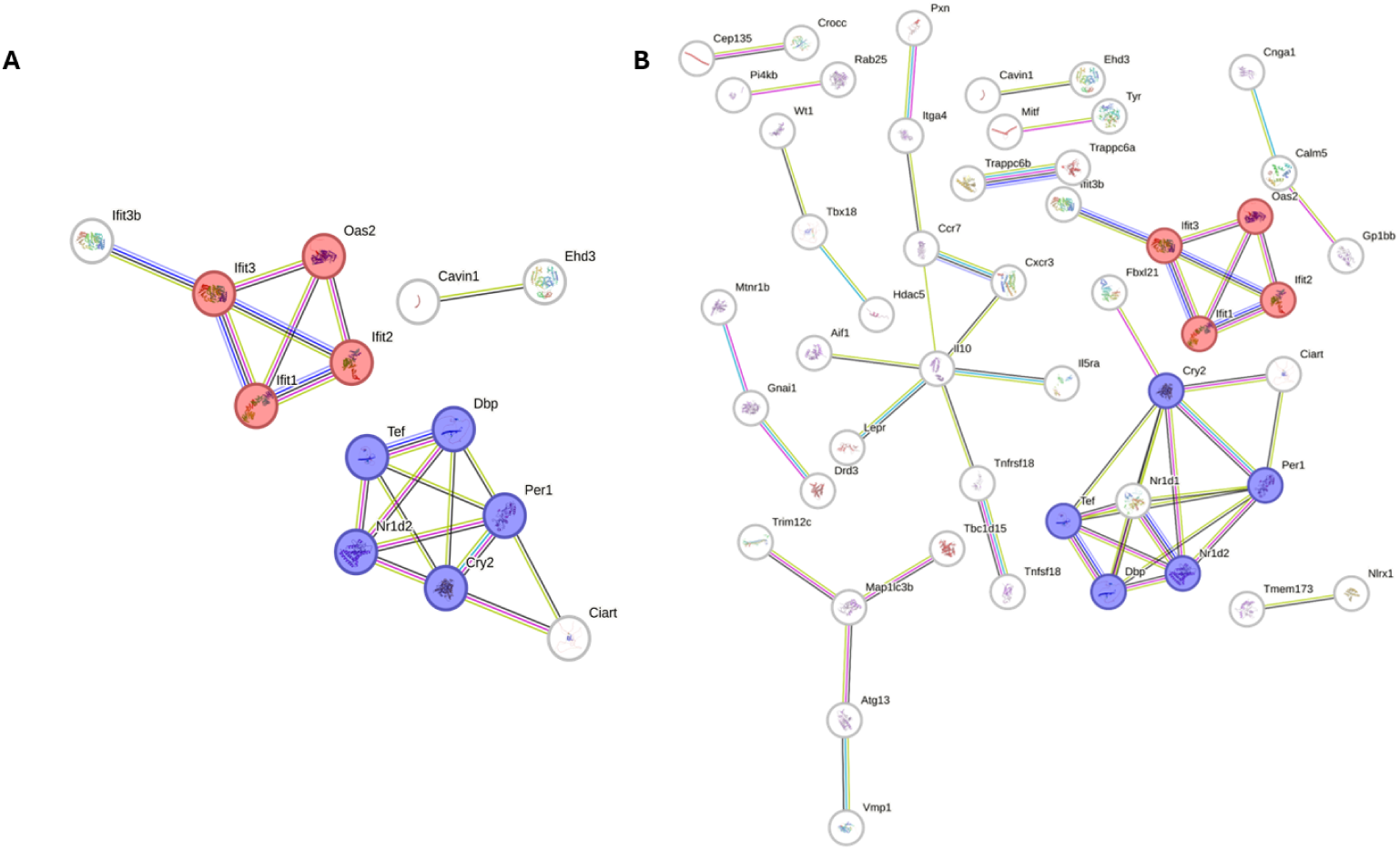
PPI networks derived from A) the 80-NN + AS model and B) the SC + WN model. These networks show a cluster containing several core clock components with only edges with a minimum interaction score of 0.7 (high confidence) included for ease of visualization. Core clock components are denoted in purple and IL-27 signaling components are denoted in red.

## 4 Discussion

Analysis of multi-omics data has long been a hurdle in the circadian field. Our goal in creating the D-CAF pipeline was to enable simultaneous co-expression analysis of circadian, or any type, of transcript and protein time series measurements. To this end, we utilized our LIMBR and ECHO software to successfully generate comparable relative expression curves for raw transcriptomic and proteomic time series. The successful integration is evidenced by the lack of clusters formed solely of isolated transcript or protein data types after unweighted analysis (Figure S1.2). Following this, co-expression analysis pathways for the multi-omic data were created using two methods: unweighted and weighted network analysis, both of which partitioned the data into densely connected clusters that were robust to Gaussian noise perturbation (Figure 4). This robustness, combined with the small deviation in rhythm parameters within each cluster, indicated this approach identified small and large groups that may have regulatory relationships, with the unweighted networks identifying small cluster groups that are highly related (Figures 5 and 7A) and the weighted networks granting insights into less related clusters that granted a more global view of the circadian network (Figure 7B). In total, the combination of clustering approaches were useful to find robust local and global relationships between known clock genes/proteins and other genes/proteins not previously found to be related to the circadian clock. With this, our D-CAF framework can be used by others to similarly analyze their own multi-omic circadian data.

As an example of these novel relationships, by unweighted analysis we found that the transcripts Ptcra and Tm9sf1, and protein NTAL, were contained in the same cluster as known clock genes Figure 5A. Ptcra regulates early T-cell development, Tm9sf1 has been shown to be involved in the inflammatory response to acute lung injury, and NTAL is important for the signaling of mast cells that control the immune system, suggesting further pathways by which circadian rhythms tune the immune system ([49], [52], [53]). Using weighted analysis, we highlighted components of the interferon (IFN) signaling pathway (Ifit1, Ifit2, Ifit3, and Oas2), which are a major subclass of IFN-stimulated genes (ISGs) known to coordinate innate immune signaling pathways and directly inhibit viral protein synthesis. ([54]). These ISGs were significantly represented (P-value = 0.0142) in a cluster along with core circadian clock components Cry2 and Per1, as well as accessory components Ciart, Nr1d1, Nr1d2, Tef, Dbp, and Hdac5.

Circadian rhythms have been implicated in ISG responses across various tissues, including the skin, lungs, and liver. In a study using pharmacological activation on murine skin, ISG expression was found to vary depending on the treatment time within the circadian cycle, primarily eliciting responses from epidermal immune cells, particularly monocytes ([55]). Additionally, systemic Bmal1 knockout mice exhibited an amplified ISG response in both skin and isolated epidermis, suggesting that mice lacking circadian rhythms may experience heightened activation of the IFN pathway. Transcriptomic data from primary human hepatocytes revealed that inflammatory ISGs exhibit circadian-regulated expression patterns. However, when Bmal1 expression is suppressed, a down-regulation of ISGs is observed ([56]). This suggests that the induction of ISGs varies depending on the time of stimulation, potentially leading to innate responses that fluctuate throughout the circadian day. SARS-CoV-2 has been demonstrated to also induce IFNs and ISGs, both in *in vivo* and *in vitro* airway mucosa ([57]; [58]). ISGs, particularly members of the Ifit family, have been shown to exhibit antiviral activity against SARS-CoV-2 ([59]). In Bmal1-silenced cells, the induction of ISGs resulted in a decreased expression of the viral receptor angiotensin-converting enzyme 2 (ACE2) and reduced viral entry into lung epithelial cells ([60]). This increase in ISG transcripts could be regulated at various stages of the IFN sensing and signaling pathways through direct or indirect mechanisms. These instances in skin, liver, and lung, along with the relationships predicted by our networks, suggest that the ISG antiviral response varies depending on the time of day.

There are limitations associated with this work. First, the network parameters were chosen based on the properties of the data specifically used in this study (i.e., 24 time points over 48 hrs), and networks for other datasets may not yield the same results. Similarly, a very conservative BH adjusted p-value threshold was used to filter the data, which may result in too many rhythms being filtered out for smaller datasets. As such, the properties of datasets should be evaluated and used when tuning the parameters used to apply the D-CAF framework.

In summary, we found D-CAF created a framework to successfully integrate multiomic data to find clusters of highly similar rhythms with biological relevance using a validated and multifaceted analysis. The pairing of unweighted and weighted network analysis facilitated the identification of relationships between clock genes/proteins and novel genes/proteins with a high correlation while simultaneously allowing for the broad analysis of the circadian network. While any model generated must be validated, D-CAF presents a novel method to perform multi-omic co-expression analysis of longitudinal omics data to determine previously unknown relationships.

## 5 Declarations

### 5.1 Ethics approval and consent to participate

Not applicable

### 5.2 Consent for publication

Not applicable

### 5.3 Availability of data and materials

The source code and data are available at https://github.com/ChuahResearchGroup/DCAF

### 5.4 Competing Interests

The authors have no competing interests to declare.

### 5.5 Funding

This work was supported by a NIH-National Institute of Aging T32 Fellowship (T32AG078123) (to J.C. and C.C.), and a NIH-National Institute of Biomedical Imaging and Bioengineering Grant (U01EB022546), a NIH-National Institute of General Medical Sciences Grant (R35GM128687), and an NSF CAREER Award 2045674 (to J.M.H.).

### 5.6 Author’ contributions

Conceptualization: J.C., J.H., J.M.H; Formal Analysis: J.C.; Funding Acquisition: J.C., C.C., J.H., J.M.H.; Investigation: J.C., J.H., J.M.H.; Methodology: J.C., J.H., J.M.H.; Software: J.C.; Visualization: J.C.; Writing-original draft: J.C., J.M.H; Writing-review and editing: J.C., C.C., J.H., and J.M.H.

## 5.7 Acknowledgments

The authors would like to thank Sharleen Buel and Meaghan Jankowski for their contributions to this study. This work was supported by a NIH-National Institute of Aging T32 Fellowship (T32AG078123) (to J.C. and C.C.), and a NIH-National Institute of Biomedical Imaging and Bioengineering Grant (U01EB022546), a NIH-National Institute of General Medical Sciences Grant (R35GM128687), and an NSF CAREER Award 2045674 (to J.M.H.).

## Notes

https://github.com/ChuahResearchGroup/DCAF

### Competing Interest Statement

The authors have declared no competing interest.

