## Supplemental File 1 for "Dual-Approach Co-expression Analysis Framework (D-CAF) Enables Identification of Novel Circadian Regulation From Multi-Omic Timeseries Data"

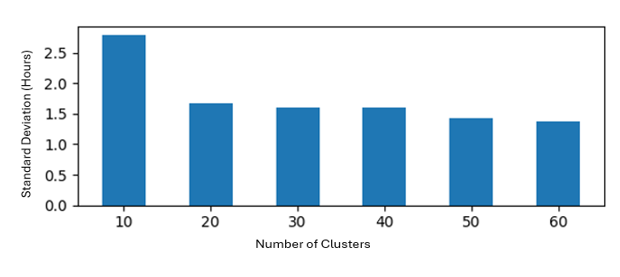

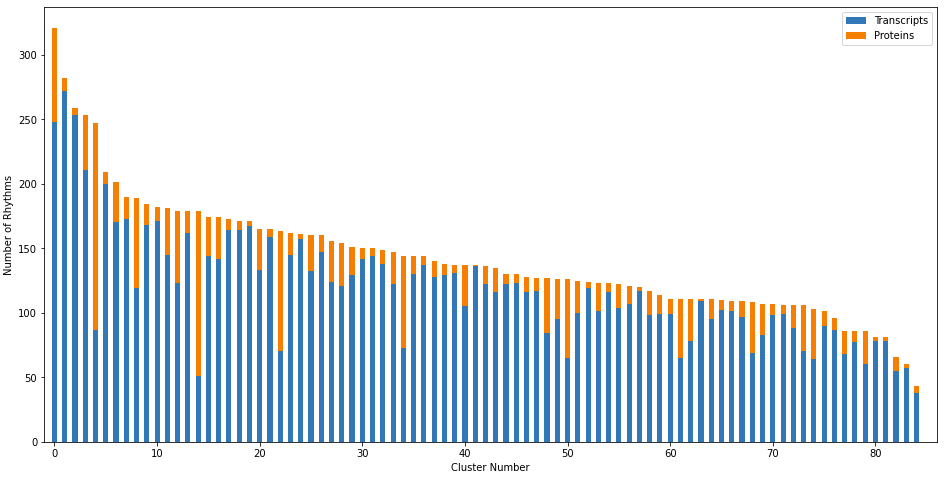
Fig S1.1: Within-cluster standard deviation of parameters based on the number of clusters from the SC + WN model. The X-axis shows the number of clusters used to construct the spectral clustering model, while the y-axis shows the standard deviation (A) shows the standard deviation of the within-cluster period, and (B) shows the standard deviation of the within-cluster phase shift.

Fig S1.2: Number of rhythms per cluster of the 80-NN + AS model. Blue regions show the number of transcripts in the cluster, and orange shows the number of proteins. In the filtered dataset, there are 10,651 transcripts and 2026 proteins

| Cluster Number (out of 84) | Standard Deviation of Cluster Rhythm Period (Hours) | Standard Deviation of Cluster Rhythm Phase Shift (Hours) |
| --- | --- | --- |
| 8 | 1.866 | 1.082 |
| 10 | 1.679 | 2.031 |
| 17 | 1.29 | 1.949 |
| 24 | 1.383 | 1.31 |
| 26 | 0.609 | 0.957 |
| 38 | 1.642 | 1.951 |
| 41 | 1.911 | 1.52 |
| 42 | 1.305 | 1.377 |
| 44 | 3.245 | 2.678 |
| 53 | 3.066 | 1.168 |
| 58 | 1.169 | 1.405 |
| 62 | 3.22 | 1.215 |
| 63 | 0.974 | 1.542 |
| 66 | 2.437 | 0.754 |
| 74 | 0.996 | 1.419 |
| 77 | 3.107 | 0.944 |
| 78 | 0.767 | 2.071 |
| 81 | 2.986 | 1.534 |
| Average | 1.870 | 1.495 |

Table S1.1: The average parameters in each circadian cluster for the 80-NN + AS model. Cluster numbers correspond to Supplemental File 2. Within-cluster standard deviations of the period and phase shift (calculated by ECHO) for each circadian cluster were computed. Smaller deviation indicates more similarity between the rhythms in a cluster.

| Cluster Number (Out of 30) | Medium Confidence P-Value | High Confidence P-Value | Very High Confidence P-Value |
| --- | --- | --- | --- |
| 11 | 5.96E-06 | 6.28E-06 | 0.0673 |
| 13 | 0.00296 | 0.00523 | 0.121 |
| 17 | 0.273 | 0.331 | 0.183 |
| 19 | 0.00697 | 0.00441 | 3.03E-04 |
| 25 | 0.167 | 0.34 | 0.265 |

Table S1.2: PPI Enrichment Interaction P-Values of 5 clusters from the Spectral clustering weighted co-expression analysis model. Levels of confidence indicate the probability that links in the StringDB database exist in the KEGG database (Medium = 0.4, High = 0.7, Very High = 0.9).
